# Sympathetic vasomotion as an early marker of hemorrhage

**DOI:** 10.1101/2025.08.23.671352

**Authors:** Peter Ricci Pellegrino, Alicia M. Schiller, Iraklis I. Pipinos, Irving H. Zucker, Han-Jun Wang

## Abstract

Each year, over 1.8 million people die from hemorrhagic shock, and, since the median time from onset to death is only two hours, early recognition is the cornerstone of management. The sympathetic nervous system is the fastest physiological hemodynamic compensatory mechanism, and we have developed a novel measure of sympathetic vascular control called sympathetic vasomotion which could serve as an early marker of hemorrhage. We performed unilateral renal denervation on six rabbits and instrumented these rabbits with bilateral renal flow probes and arterial pressure telemeters to allow for measurement of sympathetic vasomotion in paired vascular beds that differed only by sympathetic innervation. After a two-week recovery period, conscious rabbits then underwent controlled blood withdrawal via an auricular arterial catheter to simulate hemorrhage. Vasomotion differences between innervated and denervated kidneys in admittance gain, phase shift, and coherence increased significantly prior to increases in heart rate or decreases in blood pressure. These data suggest that sympathetic vasomotion could be a useful physiologically based biomarker for the early detection of hemorrhage. Further studies are needed to evaluate the utility of monitoring the sympathetic nervous system in clinical settings.

**NEW & NOTEWORTHY:** Sympathetic vasomotion, a novel marker of sympathetic outflow, increases prior to other hemodynamic changes. Sympathetic vasomotion could serve as an early detection tool for hemorrhage that facilitates prompt and precise resuscitation.

## INTRODUCTION

Hemorrhagic shock is a significant source of morbidity and mortality. Over 1.8 million people die annually from hemorrhagic shock, accounting for over 86 million years of life lost(1). The median time from bleeding onset to death is only 2 hours. Therefore, early recognition is the cornerstone of management(2).

Because of the clinical importance of the early detection of hemorrhage, several measures of hemodynamic instability have been developed(3, 5, 7, 10, 33). Some of the leading measures are widely available clinically but are based on opaque machine learning algorithms, providing clinicians with values of unclear significance and provenance, and, despite significant industry investment, the added value of these measures in pragmatic clinical trials remains unclear(4, 6, 8). A physiologically based measure to detect hemodynamic instability and facilitate anticipatory management of hemorrhage could address the shortcomings of current measures, decrease clinical uncertainty, and improve patient outcomes.

The sympathetic nervous system is the master regulator of short-term circulatory homeostasis, integrating multimodal sensory input and coordinating changes in venous capacitance, arteriolar tone, heart rate, and humoral activation to buffer a wide range of hemodynamic stressors(9). In the case of hemorrhage, low-pressure baroreceptors sense decreases in effective circulating volume, resulting in increases in sympathetic outflow that work to maintain organ perfusion(11). The role of the sympathetic nervous system as the first responder to circulatory challenges makes it an ideal physiological marker for the early detection of hemorrhage. Unfortunately, from a practical standpoint, its utility is limited by the fact that clinicians do not currently have a reliable way to measure sympathetic nerve activity.

Our recent work has identified a novel method that leverages simultaneous recordings of blood pressure and blood flow to quantify sympathetic outflow, termed sympathetic vasomotion(12, 13). By enabling clinicians to measure sympathetic vasomotion non-invasively, this technology could allow for earlier treatment and better management of hemorrhage. Our previous work with sympathetic vasomotion has shown that sympathetic vasomotion is decreased by a broad array of sympatholytic stimuli, including surgical denervation, catheter-based radiofrequency denervation, intrathecal blockade, and ganglionic blockade, but we have never tested the effect of a sympathoexcitatory stimulus, like hemorrhage, on sympathetic vasomotion.

We hypothesized that hemorrhage would increase sympathetic vasomotion. We tested this hypothesis in a graded model of hemorrhage using a conscious, chronically instrumented rabbit model to determine the sensitivity of sympathetic vasomotion to blood loss and to eliminate the potentially confounding effects of general anesthesia. These chronically instrumented rabbits underwent bilateral renal blood flow probe implantation and unilateral surgical denervation. The bilateral nature of the kidneys and the isolated autonomic innervation of this organ allows for an elegant model in which two analogous vascular beds can differ only by their innervation status without complicating animal factors like sensorimotor loss and debility.

## MATERIALS AND METHODS

All methods are further detailed in the Online Supplement.

### Data and Materials Availability

Data and source code used for this study are available on figshare (https://doi.org/10.6084/m9.figshare.29737739). The corresponding author is responsible for the maintenance of this repository.

### Rabbit Instrumentation

Experiments were carried out on adult male New Zealand White rabbits. All experiments were reviewed and approved by our Institutional Animal Care and Use Committee and carried out in accordance with the NIH Guide for the Care and Use of Laboratory Animals. Six rabbits were instrumented with arterial pressure (AP) telemeters and bilateral renal blood flow (RBF) probes and underwent unilateral surgical renal denervation under general anesthesia. This left one kidney fully innervated (INV) with the contralateral denervated (DNx) kidney exposed to the same hemodynamic input, perfusion pressure, and circulating neurohumoral environment (Figure 1). Functional renal denervation was validated by demonstrating an abrogation of the renovascular response to the nasopharyngeal reflex to cigarette smoke(14, 15).

**Figure 1.**
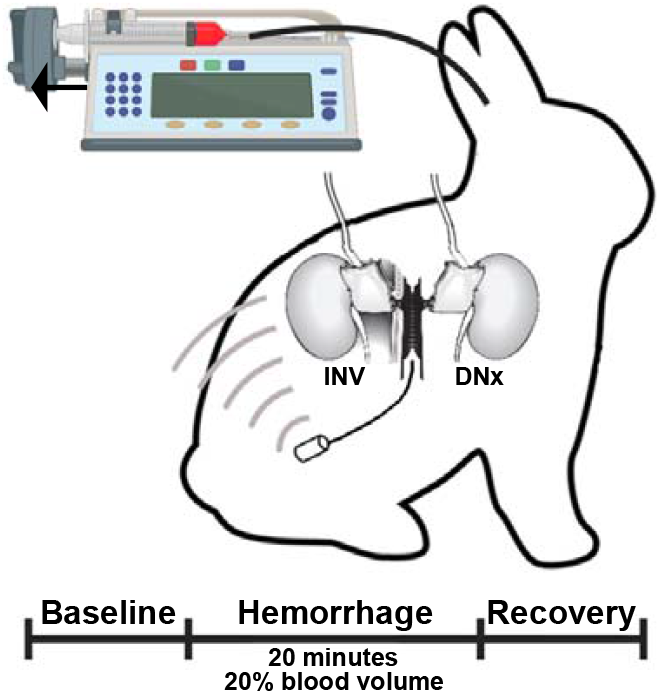
Experimental paradigm. Rabbits underwent unilateral surgical denervation and instrumentation with an arterial pressure telemeter and bilateral volumetric renal blood flow probes. At least two weeks later, rabbits underwent controlled hemorrhage via an auricular artery catheter that was connected to a syringe pump run in reverse to remove 1% estimated blood volume per minute for 20 minutes.

### Hemorrhage Experiments

After a two-week recovery period during which the rabbits were acclimated to a quiet, dimly lit procedure room, catheters were placed in the central auricular artery and the marginal ear vein under local anesthesia with 1% lidocaine. As a control, an automated syringe pump was run at a constant rate disconnected from both the intravenous and intraarterial catheters for at least 15 minutes; the last 10 minutes of data from this were used as baseline data. The auricular arterial catheter was then connected to a syringe containing heparin that was loaded on the automated pump in reverse, resulting in a constant removal of arterial blood. This setup was used to remove 1% of the estimated blood volume (60 mL/kg) per minute for 20 minutes, resulting in removal of 20% of the estimated blood volume to mimic Class II normotensive shock. The syringe pump was then stopped and another 10 minutes of data were acquired and are reported as the recovery period. At the end of the recovery period, data acquisition was terminated, and the withdrawn blood was administered via intermittent intravenous boluses.

### Data Analysis

Sympathetic vasomotion analysis was performed as described previously(12, 13). In brief, a time-varying two-element Windkessel model of the renal circulation was used to extract the beat-to-beat resistive RBF time series, which was in turn used to calculate the AP-resistive RBF time-varying transfer function through bispectral wavelet analysis. Based on the physical relationship between pressure and flow, the three components of the AP-resistive RBF time-varying transfer function each have a unique physiological interpretation. Specifically, admittance gain reflects the magnitude of transduction of pressure into flow with lower admittance gain reflecting active vasoconstrictive behavior. Phase shift indicates timing differences between oscillations in pressure and flow, with autoregulatory mechanisms creating a positive phase shift, baroreflex-mediated vascular control resulting in negative phase shift, and passive Poiseuille flow resulting in zero phase shift. Coherence reflects the consistency of the pressure-flow relationship over time and is thus inversely related to the amount of active vascular control. Occurrence histograms of admittance gain, phase shift, and coherence were calculated for each kidney in the baseline state as well as over four different time periods during hemorrhage (0-5 minutes, 5-10 minutes, 10-15 minutes, and 15-20 minutes) and in the recovery state. These individual occurrence histograms were used for subsequent group analysis, and the mean group data are displayed as contour plots. Vasomotion difference maps between the INV and DNx kidneys were computed in two ways. The first set calculated t-statistics for each independent variable pair (e.g., frequency-admittance gain bin, frequency-phase shift bin) using two-tailed, paired t-tests intended to convey directionality, magnitude, and consistency of differences but not statistical significance per se. The second set of difference maps were calculated as the occurrence difference between INV and DNx kidneys. Both were used to calculate the sympathetic vasomotion magnitude for admittance gain, phase shift, and coherence. This was performed by summing the total occurrence difference between INV and DNx kidneys at each bin where |t| > 2.571 (P < 0.05 for n = 6) in the t-statistic difference map. Total renal sympathetic vasomotion magnitude was calculated as the square root of the sum of squares for the admittance gain, phase shift, and coherence.

### Statistical Testing

All data are displayed as mean ± the standard error of the mean. Statistical testing of simple, normally distributed measures (e.g., arterial pressure, heart rate, renal blood flow, and sympathetic vasomotion magnitude) was conducted as a repeated measures ANOVA with appropriate within-subjects factors (e.g., state, innervation status) with Greenhouse-Geisser correction for sphericity and α = 0.05. Post-hoc statistical testing versus the baseline state was performed, and the Holm-Bonferroni method for multiple comparisons was used. Consistent with our prior work, for cumulative sympathetic vasomotion, we also performed non-parametric cumulative difference mass testing using t-statistic maps with a threshold of 2.571, which allows for a priori significance testing of non-independent multidimensional data while addressing the multiple comparisons issues with such data(16). The shuffled null data are displayed as violin plots alongside the actual cumulative t-statistics along with the non-parametric P values.

## RESULTS

### Nasopharyngeal Reflex

Functional renal denervation was verified by the nasopharyngeal reflex(17). Perinasal administration of a noxious stimulus in rabbits results in increases in both parasympathetic and sympathetic outflow, manifesting as bradycardia and a sympathetically mediated decrease in renal blood flow in INV kidneys (Figure S1). This renal vasoconstrictive response is absent in DNx kidneys.

### Hemodynamics

Hemodynamics for the hemorrhage experiment are shown in Figure 2. Hemorrhage resulted in a statistically significant decrease in renal blood flow (P < 0.001), and there was a statistically significant interaction between hemorrhage and innervation status (P < 0.001) with blood flow decreasing more in DNx kidneys than their INV counterparts. While this reached statistical significance, it should be noted that the magnitude of this difference (7 mL/min decrease for INV vs. 8 mL/min for DNx, respectively) is physiologically insignificant. Heart rate increased significantly (P < 0.001) although not until 10-15 minutes of hemorrhage. Mean arterial pressure did not significantly change over the course of the experiment, consistent with the targeted normotensive (Class II) hemorrhage.

**Figure 2.**
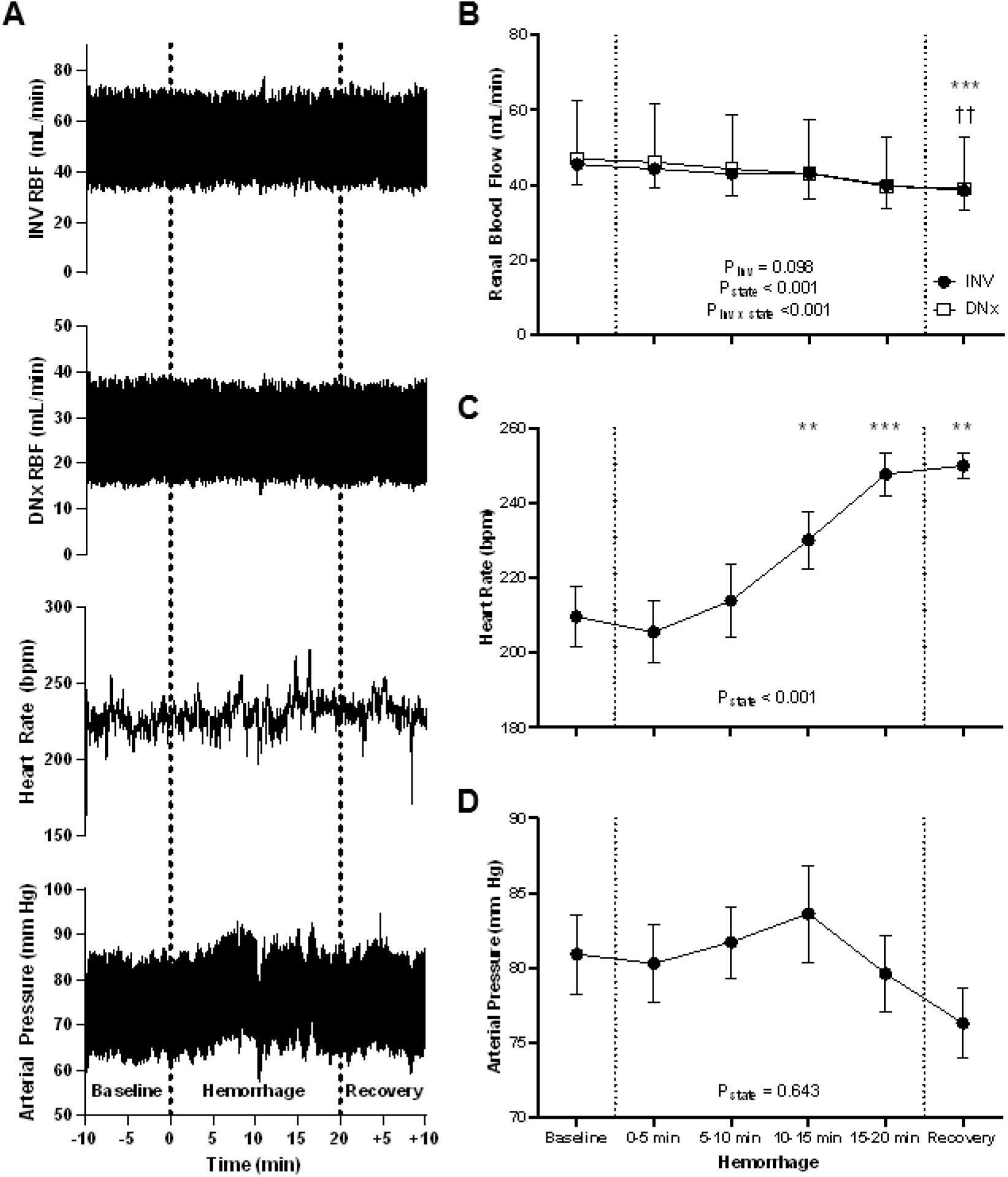
Hemodynamics. (A) Hemodynamic parameters, including renal blood flow of the innervated (INV) and denervated (DNx) kidneys, heart rate, and arterial pressure for one representative rabbit. (B) Mean renal blood flow decreased over the course of the experiment for both INV and DNx kidneys with a statistically greater change in DNx kidneys. (C) Heart rate increased significantly with hemorrhage, but not until 10-15 minutes of blood withdrawal. (D) Mean arterial pressure did not change significantly over the course of the experiment. Statistical testing was performed via repeated measures ANOVA with one (C, D) or two (B) factors, and post-hoc testing versus baseline was performed with Holm-Bonferroni-corrected paired t-tests. n = 6. * P < 0.05, ** P < 0.01, *** P < 0.005 vs. baseline, †† P < 0.01 vs. baseline for DNx. RBF, renal blood flow.

### Admittance Gain

Admittance gain quantifies the magnitude of the transduction of arterial pressure oscillations into renal blood flow with respect to passive Poiseuille flow. Lower admittance gain indicates active buffering of arterial pressure oscillations by vasoconstrictive or autoregulatory mechanisms. At baseline, there was more low admittance gain behavior in INV kidneys and more high admittance gain behavior in DNx kidneys as would be expected due to sympathetic vasoconstriction (Figure 3A, Figure S2). These differences increased with hemorrhage, reaching statistical significance at 0-5 minutes of hemorrhage and peaking at 5-10 minutes of hemorrhage (Figure 3B-D).

**Figure 3.**
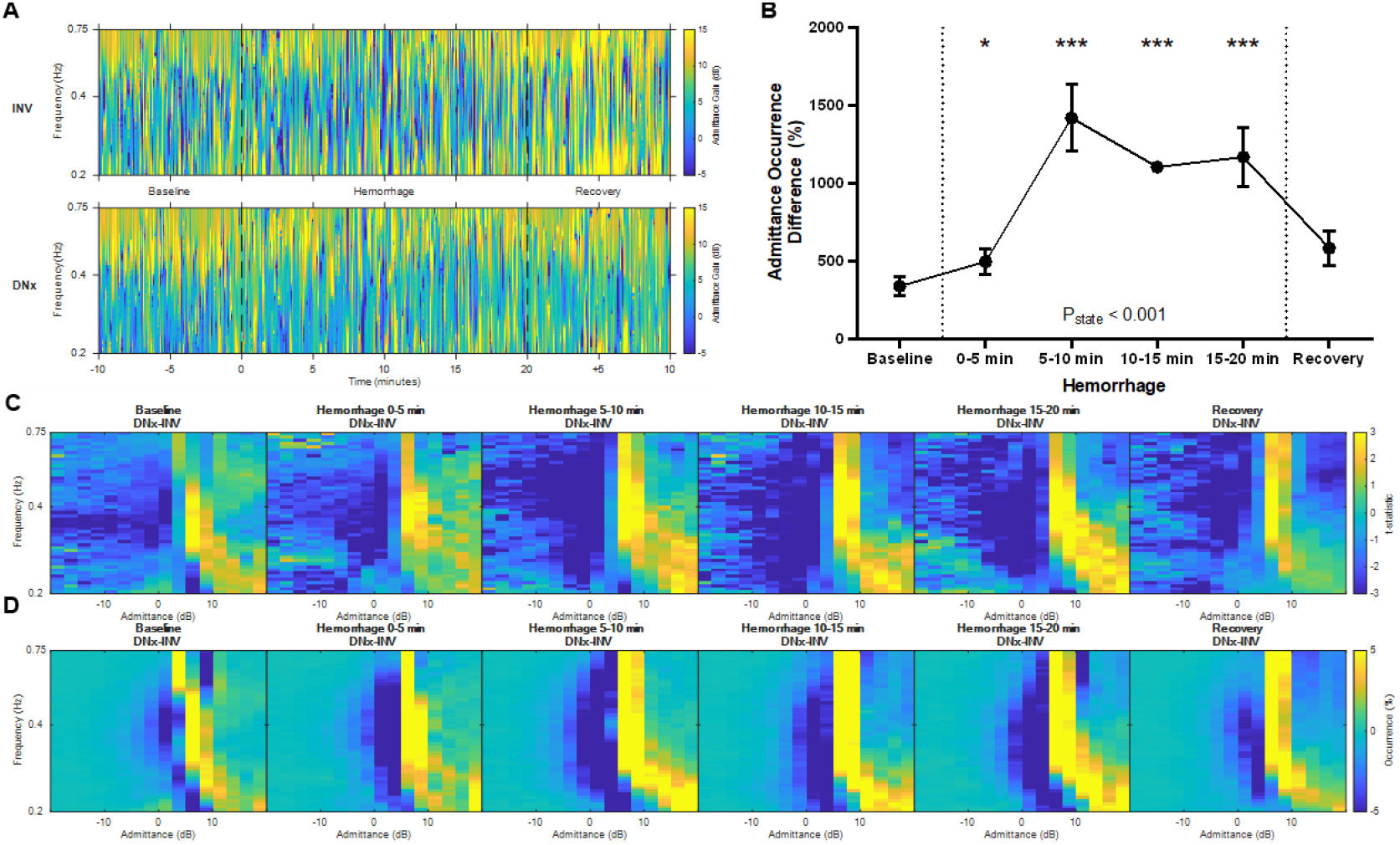
Admittance Gain. (A) Representative time-frequency plots showing admittance gain of the innervated (INV) and denervated (DNx) kidneys for one representative rabbit. (B) Admittance gain sympathetic vasomotion magnitude increased with hemorrhage. (C) Vasomotion difference maps of t-statistics between INV and DNx kidneys and (D) vasomotion difference maps of mean occurrence between INV and DNx kidneys show increases in low admittance gain behavior with hemorrhage. Statistical testing was performed on the data in B using a repeated measures ANOVA with post-hoc testing versus baseline using Holm-Bonferroni-corrected paired t-tests n = 6. * P < 0.05, ** P < 0.01, *** P < 0.005 vs. baseline.

### Phase Shift

Phase shift quantifies the temporal relationship between oscillations in blood pressure and blood flow and provides insight into the directionality of vascular control mechanisms. Passive Poiseuille flow is characterized by synchrony between blood pressure and flow and thus a phase shift of zero. Negative phase shift behavior indicates that arterial pressure oscillations precede renovascular modulation, consistent with baroreflex-mediated vascular control, while positive phase shift behavior indicates that blood flow precedes renovascular modulation, consistent with autoregulatory vascular control. At baseline, there was more negative phase shift behavior in INV kidneys and more zero and positive phase shift behavior in DNx kidneys (Figure 4A, Figure S3). Hemorrhage significantly increased these differences, reaching statistical significance at 5-10 minutes of hemorrhage and remaining statistically significant for the remainder of the experiment (Figure 4B-D).

**Figure 4.**
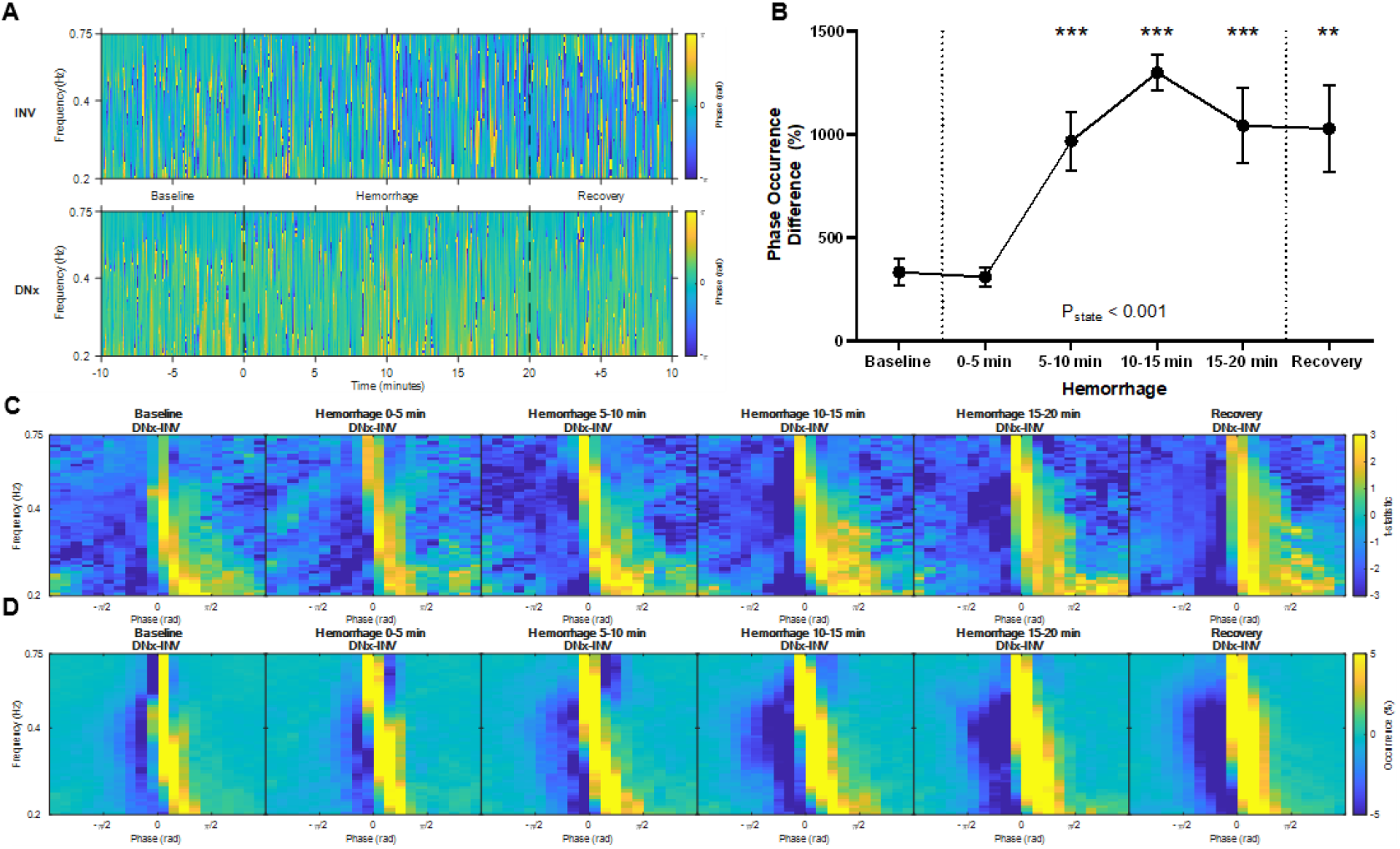
Phase Shift. (A) Representative time-frequency plots showing phase shift of the innervated (INV) and denervated (DNx) kidneys for one representative rabbit. (B) Phase shift sympathetic vasomotion magnitude increased with hemorrhage. (C) Vasomotion difference maps of t-statistics between INV and DNx kidneys and (D) vasomotion difference maps of mean occurrence between INV and DNx kidneys show increases in negative phase behavior with hemorrhage. Statistical testing was performed on the data in B using a repeated measures ANOVA with post-hoc testing versus baseline using Holm-Bonferroni-corrected paired t-tests. n = 6. * P < 0.05, ** P < 0.01, *** P < 0.005 vs. baseline.

### Coherence

Coherence quantifies the consistency in the pressure-flow relationship over time and correlates inversely with active vascular modulation. Surgical denervation removes an active vascular control mechanism, and, accordingly, DNx kidneys displayed more high coherence behavior than INV kidneys (Figure 5A, Figure S4). These differences increased with hemorrhage, reaching statistical significance in the first five minutes of hemorrhage and remaining elevated for the remainder of the experiment.

**Figure 5.**
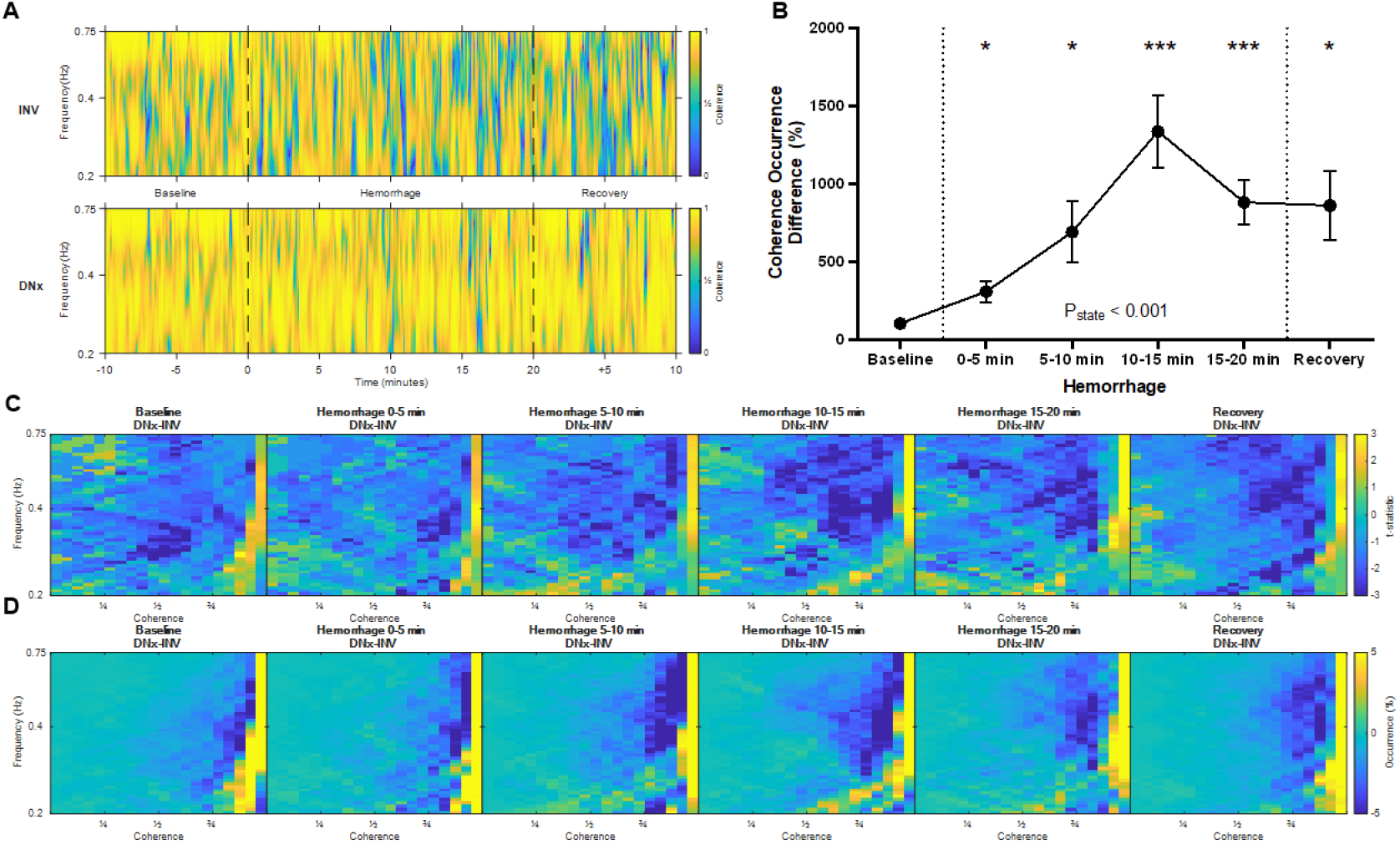
Coherence. (A) Representative time-frequency plots showing coherence of the innervated (INV) and denervated (DNx) kidneys for one representative rabbit. (B) Coherence sympathetic vasomotion magnitude increased with hemorrhage. (C) Vasomotion difference maps of t-statistics between INV and DNx kidneys and (D) vasomotion difference maps of mean occurrence between INV and DNx kidneys show increases in low coherence behavior with hemorrhage. Statistical testing was performed on the data in B using a repeated measures ANOVA with post-hoc testing versus baseline using Holm-Bonferroni-corrected paired t-tests. n = 6. * P < 0.05, ** P < 0.01, *** P < 0.005 vs. baseline.

### Total Sympathetic Vasomotion

The data from admittance gain, phase shift, and coherence of the arterial pressure-resistive blood flow time-varying transfer function were aggregated in two measures: total renal sympathetic vasomotion magnitude and the total vasomotion profile difference. Total renal sympathetic vasomotion, the square root of the sum of squares for the transfer function components, is displayed in Figure 6A. Total renal sympathetic vasomotion difference increases in the first five minutes of hemorrhage and remains statistically higher throughout the remainder of the experiment. The total vasomotion profile difference is a statistical group measure that reflects the statistical differences across all components of the time-varying pressure-resistive flow transfer function (Figure 6B). The total vasomotion profile difference was statistically significant at baseline, and the magnitude of this difference increased with hemorrhage, peaking between 10-15 minutes of hemorrhage. The difference between INV and DNx kidneys was statistically significant for all states (from baseline to recovery).

**Figure 6.**
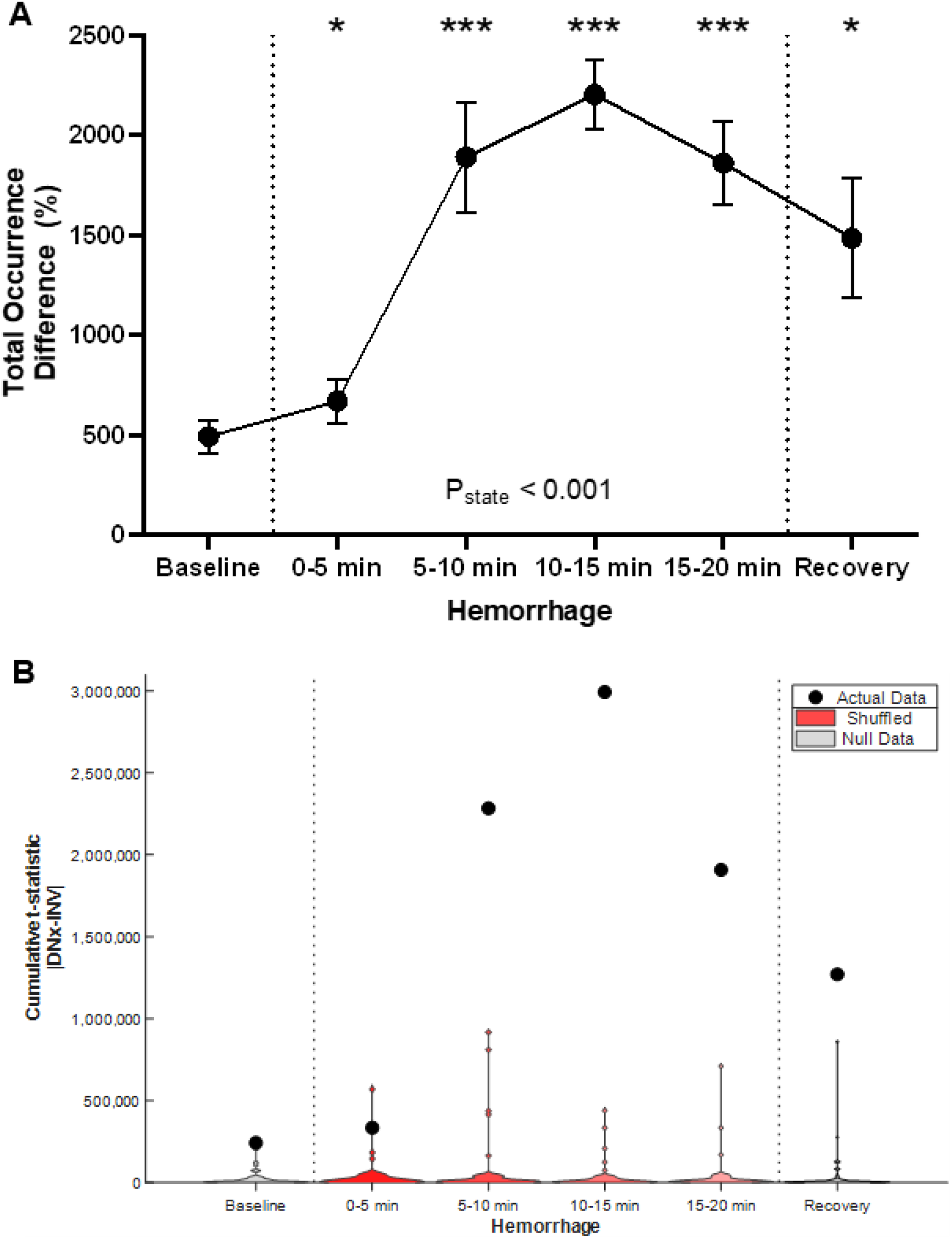
Total Vasomotion. Hemorrhage increased (A) total renal sympathetic vasomotion magnitude and (B) the total vasomotion profile difference. Statistical testing was performed on the data in A using a repeated measures ANOVA with post-hoc testing versus baseline using Holm-Bonferroni-corrected paired t-tests. Statistical testing was performed on the data in B using non-parametric cumulative difference mass testing using t-statistic maps with a threshold of 2.571. n = 6. * P < 0.05, ** P < 0.01, *** P < 0.005 vs. baseline. n = 6. * P < 0.05, ** P < 0.01, *** P < 0.005 vs. baseline.

## DISCUSSION

These data show that renal sympathetic vasomotion increases in response to hemorrhage. Moreover, increases in sympathetic vasomotion occur prior to increases in heart rate or decreases in blood pressure. Sympathetic vasomotion remains elevated over the course of hemorrhage. These data are consistent with the physiological role of the sympathetic nervous system as the first responder to hemodynamic stress and indicate that sympathetic vasomotion may be a useful early marker of hemorrhage.

Some of the nuances in the presented data are best interpreted after a discussion of the neural mechanisms underlying the sympathetic response to hemorrhage. Hemorrhage initially decreases effective circulating volume, which is sensed by the low-pressure baroreceptors in the atria, ventricles, and pulmonary circulation, and eventually decreases cardiac output, which is sensed by arterial baroreceptors in the aorta and carotid sinus. These baroreceptors transmit this information to key autonomic nuclei like the nucleus tractus solitarius, rostral ventrolateral medulla, and the paraventricular nucleus of the hypothalamus(18, 19). Integration occurs at these key autonomic nuclei with distant modulation from higher-order structures, including the thalamus, the insular cortices, the anterior cingulate gyrus, and the orbitofrontal cortex of the cortical autonomic network(20). This results in increased sympathetic outflow, mobilizing blood volume via venoconstriction, constricting arterioles to maintain perfusion pressure, and increasing heart rate.

The time delay between the activation of low-pressure and high-pressure baroreceptors may explain why admittance gain and coherence differences increase within the first five minutes of hemorrhage while phase shift differences did not increase until 5-10 minutes of hemorrhage. Admittance gain and coherence measure active vasoconstriction and vascular control, which would be increased by unloading of the low-pressure baroreceptors. For phase shift, the observed increases are instead indicative of increased baroreflex-mediated renovascular control, which would not occur until the arterial baroreceptors are unloaded later in hemorrhage. Additionally, admittance gain peaked at 5-10 minutes of hemorrhage and was no longer statistically different from baseline during the recovery phase (P = 0.07). Because of how admittance gain is normalized to mean renovascular conductance, this measure likely reflects the rate of change in sympathetically mediated vascular tone rather than the sympathetic vascular tone per se. Similarly, conflicting neural reflexes may underpin the delayed tachycardia despite clear increases in renal sympathetic tone. Decreases in venous return due to hemorrhage may unload the atrial receptors and inhibit the Bainbridge reflex, which would oppose the expected tachycardia from increases in sympathetic outflow. Subsequent experiments involving measurement of cardiac filling pressures and cardiac output and manipulation of the neural circuitry involved in the response to hemorrhage, including sinoaortic denervation, could better characterize the mechanism of these differences.

The renal circulation was used for these experiments for two reasons. First, as stated previously, as a bilateral visceral organ, the kidneys offer an elegant model where two circulatory beds can differ strictly by innervation status while being exposed to the same systemic stimulus, perfusion pressure, and circulating humoral milieu. Second, the renal circulation and its innervation are of significant clinical interest. In the inpatient setting, renal circulatory changes are a driving factor in the most common causes of acute kidney injury, including sepsis and hemorrhage (21, 22). On a chronic basis, hypertension and diabetes profoundly impact the renal circulation on a structural and functional level, and, in doing so, lead to renal dysfunction(23). Moreover, elevated renal sympathetic outflow plays a causal role in human hypertension, which underpins the minimally invasive renal denervation technologies employed clinically, and in the renal dysfunction seen in chronic heart failure (24–28).

Our finding that renal sympathetic vasomotion is an early marker of hemorrhage highlights the broader principle that sympathetic control of other vascular beds could also provide distinct, valuable sensors for hemodynamic stress. This first responder role of the sympathetic nervous system is likely just as strong and easier to access in peripheral tissues due to their proximity to the skin. For example, cutaneous blood flow is under tonic sympathetic inhibition and would be expected to exhibit intense vasomotion at the earliest onset of hemorrhage as the body shunts blood centrally(29). A similar early response may be measured in the limb vasculature, where sympathetic vasomotion could be measured in the femoral, brachial, or radial arteries. In contrast, the cerebral circulation, at the level of the carotid or middle cerebral artery, may display even stronger autoregulatory mechanisms than those observed in the kidney(30). Observing diverging patterns of vasomotion across different beds could provide individualized signatures for different shock states and hemodynamic stressors, providing clinicians with more useful information.

The ability to predict impending hemodynamic collapse is critical in the perioperative, critical care, and emergency settings, and other measures have been developed for this purpose. The most widely available measure in the clinical environment is the hypotension prediction index, which is based on proprietary algorithms derived from machine learning and is integrated into Acumen patient monitoring equipment (Becton, Dickinson and Company, Franklin Lakes, NJ, USA). This index has shown mixed results in clinical practice and has recently been criticized as a surrogate for mean arterial pressure(4, 6, 31). Another measure in development is the cardiac reserve index, which is also a measure that was derived from machine learning analysis of plethysmography data and has shown promise in experimental and early clinical work(32, 33). Both measures are based on opaque artificial neural networks trained on rudimentary features from subject waveforms and are not informed by autonomic neural networks or vascular physiology. While many other such measures have been proposed, sympathetic vasomotion is unique in that it is the only measure that is derived from the known physiological role of the sympathetic nervous system as the body’s first responder to hemodynamic stress.

Our study has several limitations. We did not monitor the rabbits while we retransfused the withdrawn blood, which would have provided evidence about the utility of sympathetic vasomotion as an endpoint for resuscitation. Additionally, while our work established the sensitivity of renal sympathetic vasomotion in the detection of hemorrhage, we did not test the specificity of sympathetic vasomotion for hemorrhage compared to other sympathoexcitatory stimuli. This is important in clinical contexts where pain and other modes of shock (e.g., cardiogenic, obstructive) may also increase sympathetic tone. Furthermore, as detailed previously, other circulations beyond the renal circulation may be important and more accessible sources for clinical hemodynamic monitoring. Finally, frequency-based measures from rabbits, which have physiological (e.g., cardiac, respiratory, and sympathetic) frequencies that are quite different from humans, must be reproduced in large animals prior to translating them to the clinic.

In rabbits, hemorrhage increases renal sympathetic vasomotion prior to changes in heart rate or blood pressure, which is consistent with the physiological role of the sympathetic nervous system as the first responder to hemodynamic stress. Sympathetic vasomotion could form the basis for the novel physiologically based marker of hemodynamic instability, helping clinicians recognize hemorrhage earlier and optimize resuscitation. Subsequent work will focus on translation of this work to large animals and humans.

## DATA AVAILABILITY

Data and MATLAB source code are openly available on figshare (https://doi.org/10.6084/m9.figshare.29737739).

## SUPPLEMENTAL MATERIAL

Supplemental Figs. S1-S4: https://doi.org/10.6084/m9.figshare.29941676

## ACKNOWLEDGMENTS

Preprint is available on bioRxiv.

## GRANTS

National Institutes of Health, National Institute on Aging, AG077803, and Department of Veterans Affairs, RX004632 to IIP.

National Institutes of Health, National Heart Lung and Blood Institute, HL153176 and HL172029 and Theodore Hubbard Foundation to IHZ.

National Institutes of Health, National Heart Lung and Blood Institute, HL171602, HL169205, HL152160, HL172029, and HL170127 to HJW.

## AUTHOR CONTRIBUTIONS

PRP conceived and designed the research, performed experiments, analyzed data, interpreted results of experiments, prepared figures, drafted manuscript, edited and revised manuscript, and approved final version of manuscript.

AMS conceived and designed the research, performed experiments, interpreted results of experiments, edited and revised manuscript, and approved final version of manuscript.

IIP interpreted results of experiments, edited and revised manuscript, and approved final version of manuscript.

IHZ conceived and designed the research, interpreted results of experiments, edited and revised manuscript, and approved final version of manuscript.

HJW conceived and designed the research, interpreted results of experiments, edited and revised manuscript, and approved final version of manuscript.

