## Supplemental Material for "Sympathetic vasomotion as an early marker of hemorrhage"

**Authors:** Peter Ricci Pellegrino M.D., Ph.D.[1]*, Alicia M. Schiller, Ph.D.[1], Iraklis I. Pipinos, M.D.[2], Irving H. Zucker, Ph.D.[3], Han-Jun Wang, M.D.[1]

**Affiliations:**

[1]Department of Anesthesiology, University of Nebraska Medical Center, Omaha, NE, USA

[2]Department of Surgery, University of Nebraska Medical Center, Omaha, NE, USA

[3] Department of Cellular and Integrative Physiology, University of Nebraska Medical Center, Omaha, NE

*

**Online Methods**

*Rabbits*

Experiments were carried out on six male New Zealand White rabbits ranging in weight from 3.5 to 4.2 kg (Charles River Laboratories, International, Wilmington, MA). Male animals were used to decrease heterogeneity in the data which may negatively impact the generalizability of these studies to female patients. Rabbits were housed in individual cages in a temperature-controlled room (23º C) with a 12:12 hour light-dark cycle (lights on at 07:00, lights off at 19:00). Rabbits were given *ad libitum* access to reverse-osmosis purified water and high-fiber rabbit chow (Teklad 2031, Envigo RMS, Indianapolis, IN, USA). All experiments were reviewed and approved by the University of Nebraska Medical Center Institutional Animal Care and Use Committee and carried out in accordance with the Guide for the Care and Use of Laboratory Animals of the National Institutes of Health.

*Chronic Instrumentation of Rabbits*

Six rabbits were instrumented with arterial pressure (AP) telemeters and bilateral renal blood flow (RBF) probes and underwent unilateral surgical renal denervation. Rabbits were anesthetized using ketamine (17.5 mg/kg intramuscular) and xylazine (2.9 mg/kg intramuscular) for induction and isoflurane (1-4% via endotracheal tube) for maintenance and mechanically ventilated during surgery. After a femoral cutdown was performed, the pressure transduction catheter of a pressure telemeter (PA-C40, Data Sciences International, St. Paul, MN) was inserted into the abdominal aorta via the right femoral artery and the telemeter was implanted in a pocket through the same incision in the ipsilateral inguinal area. Both the catheter and telemeter were secured with non-absorbable braided suture. Kidneys were randomized to denervation using a randomization sequence from Random.org. The kidney on the side randomized for denervation was then approached via a flank incision, and the renal arteries and veins were carefully dissected using glass rods. The renal artery was stripped of all visible neural tissue under a surgical microscope. A 2-mm renal flow probe (Transonic Systems, Inc., Ithaca, NY, USA) was secured around each renal artery using a piece of Silastic and 4-0 non-absorbable suture. The probe cable was tunneled beneath the skin and exited in the periscapular region of each rabbit where it was secured via a skin button. The flank incision was closed in layers. The same procedure, excluding the stripping of neural tissue under the surgical microscope, was performed on the contralateral side. The randomization was such that four rabbits underwent right renal denervation and two underwent left renal denervation in this cohort. On the day of surgery, rabbits received pre-surgical enrofloxacin (22.7 mg subcutaneous, Bayer HealthCare LLC, Shawnee Mission, KS) and buprenorphine (0.02 mg/kg subcutaneous) and a 72-hour fentanyl patch (25 mcg/hr transdermal) for post-operative analgesia. Rabbits were allowed to recover 14 days from surgery prior to data collection, during which time they were acclimated to resting quietly in a Plexiglas box in the experiment room.

*Nasopharyngeal Reflex*

After at least two weeks of recovery from instrumentation surgery, rabbits underwent functional validation of renal denervation by elicitation of the nasopharyngeal reflex. Rabbits were allowed to rest in a Plexiglas box in a dimly lit procedure room for at least 10 minutes. Pulsatile AP and bilateral RBF was digitized at a sampling frequency of 1 kHz (PowerLab 16/35, ADInstruments, Inc., Colorado Springs, CO, USA). The nasopharyngeal reflex was elicited by drawing thick cigarette smoke through its filter into a 60-mL syringe such that the entire syringe was opaque with smoke. The smoke was then gently delivered to the rabbit's face over 5-10 seconds, leading to high sympathetic and parasympathetic activation. The renal vasoconstrictive response was used as a check variable to validate the completeness of renal denervation and to ensure that the intact kidney remained innervated after its renal arterial instrumentation. We have previously documented that this surgical renal denervation nearly eliminates renal cortical norepinephrine.

This was a predefined check variable and exclusion criterion. If the DNx kidney exhibited a decrease in RVC by greater than 30%, the kidney was insufficiently denervated, and this rabbit was to be excluded from subsequent analysis. Conversely, if an innervated kidney failed to decrease RBF below 2 mL/min after nasopharyngeal reflex elicitation, this would indicate that the kidney was partially functionally denervated potentially due to dissection for flow probe instrumentation. All rabbits demonstrated functional innervation as expected according to surgical treatment and thus none were excluded.

*Hemorrhage*

On a subsequent day, the rabbits were again brought into the procedural room and placed in a Plexiglas box. The skin near the marginal ear vein and central auricular artery was sterilized with alcohol and anesthetized with infiltration of 1% lidocaine. A 22G intravenous catheter was placed into the marginal ear vein, and a 22G arterial catheter was placed into the central auricular artery; both were secured with tape. Rabbits were allowed to rest for at least 10 minutes after catheter placement. Then, as above, AP and bilateral RBF were recorded and digitized with a disconnected syringe pump running to provide the same ambient noise as during blood withdrawal. After at least ten minutes of artifact-free baseline data was acquired, the arterial catheter was connected to a 60 mL syringe loaded with 200 units of heparin, a dose chosen to prevent clotting with the syringe that would not produce a physiological anticoagulant effect when administered systemically. The heparin loaded syringe was then loaded to a syringe pump (Model '22', Harvard Apparatus, Holliston, MA, USA), which was run in reverse, retracting the syringe plunger and resulting in a constant removal of 1% of the estimated blood volume per minute. Estimated blood volume was calculated as 60 mL/kg; the syringe pump removed 0.6 mL/kg/min. The blood removal was performed for

20 minutes, resulting in removal of a total of 12 mL/kg (42 to 50.4 mL) of blood. After 20 minutes, the syringe was disconnected from the syringe pump, which continued to run disconnected from the rabbit. After 10 minutes of post-hemorrhage data collection, recording was stopped. The removed blood was then injected slowly into the intravenous catheter after flushing the catheter with saline to confirm patency. The arterial and venous catheters were removed and pressure was maintained on the arterial puncture sites for 5 minutes to reduce hematoma formation risk. Rabbits were observed for 10 minutes and then returned to their home cages and checked on later that day. There were no complications apart from minor hematoma formation from the experiment.

*Arterial Pressure Correction and Euthanasia*

On a subsequent day, the rabbits were anesthetized with ketamine (35 mg/kg) and xylazine (5.8 mg/kg) intramuscular injections and local femoral infiltrations of bupivacaine. The left femoral artery was then cannulated with a pressure transducer (Mikro-Tip, Millar, Pearland, TX), which was advanced to the abdominal aorta. The difference in the mean pressure between the acutely placed pressure transducer and the chronically implanted telemetry was used to correct for any drift in AP over time from the previously collected conscious data. Rabbits were then euthanized with a 3 mL Fatal-Plus (Vortech Pharmaceuticals, Dearborn Michigan) IV bolus after which a bilateral thoracotomy was performed.

*Data Analysis*

Sympathetic vasomotion analysis, which models a regional vascular bed with a two-component, time-varying Windkessel model and then constructs a time-varying pressure-resistive blood flow transfer function using cross-spectral analysis, was performed as described previously (doi: 10.1161/HYPERTENSIONAHA.120.15325). Analysis was performed by an investigator who was not blinded to kidney treatment modality. Signal processing of arterial blood flow and bilateral renal blood flow data was performed in MATLAB (Mathworks, Natick, MA, USA). The MATLAB code for the analysis of the data, the data, and instructions for reproduction of the figures in the manuscript are openly available on figshare. The experiment was broken down into six states: baseline (pre-hemorrhage), 0-5 minutes of hemorrhage, 5-10 minutes of hemorrhage, 10-15 minutes of hemorrhage, 15-20 minutes of hemorrhage, and recovery (post-hemorrhage, pre-transfusion of removed blood).

Cross-correlation analysis was used to synchronize the AP and RBF signals, eliminating time delays that arise from differential time delays between the telemetry-transmitted AP signals and the hard-wired RBF signals. Cardiac cycle identification was performed manually using the arterial pressure waveform. Nonlinearly sampled AP and resistive RBF time series were derived from this data using the two-component Windkessel model. These time series were then linearly resampled at 10 Hz. Wavelet analysis was performed using a continuous complex Morlet waveform with 51 logarithmically spaced scales corresponding to frequencies from 0.2 to 0.75 Hz where the dominant sympathetic vasomotor rhythms operate in the rabbit. The AP-resistive RBF wavelet cross-spectrum was divided by the AP autospectrum to yield the time-varying AP-resistive RBF transfer function. Admittance gain was expressed by convention in decibels and thus was calculated by taking the base-10 logarithm of the magnitude of the transfer function after normalization by the mean time-varying renal vascular conductance for that time segment and multiplying by 20. Phase shift was calculated as the angle of the real and imaginary components of the transfer function. Coherence was calculated in a scale-dependent manner using a time window of three scale lengths. This time-varying transfer function data was then put into group occurrence histograms. For admittance gain, histograms were constructed from 16 bins of 2.5-dB width across the range [-20, 20] for each scale. For phase shift, histograms were constructed from 20 bins of $\pi/10$ rad width across the range $[-\pi, \pi]$ for each scale. For coherence, histograms were constructed from 20 bins of 0.05 width across the range [0,1] for each scale. Edge-affected data were excluded from the occurrence histograms.

Renal sympathetic vasomotion magnitude was calculated to quantify the magnitude of renal sympathetic vascular control over the course of the experiment as the difference in vasomotion between INV and DNx kidneys. Paired t-tests were performed on occurrence histograms for admittance gain, phase shift, and coherence between INV and DNx kidneys. These t-statistic difference maps were thresholded at $|t| > 2.571$, which corresponds to a $P < 0.05$ for $n = 6$, and the raw occurrence difference between the INV and DNx kidneys was summed to calculate the renal sympathetic vasomotion magnitude for each rabbit for each transfer function component (admittance gain, phase shift, coherence) in each state (baseline, hemorrhage. The total renal sympathetic vasomotion magnitude was calculated as the square root of the sum of squares for the renal sympathetic vasomotion for admittance gain, phase shift, and coherence.

*Statistical Analysis*

All data are displayed as mean ± the standard error of the mean. All analysis was carried out in a nonblinded manner. Some data is visualized in 3D plots with t-statistics computed for each independent variable pair (e.g., frequency-admittance gain bin) in order to convey directionality, magnitude, and consistency of differences but not statistical significance per se. These t-statistics were calculated using a two-tailed, paired t-test. Statistical testing of hemodynamic data and renal sympathetic vasomotion magnitude was conducted with repeated measures analysis of variance with appropriate within-subjects factors (e.g., state, innervation) and $\alpha = 0.05$ in SPSS (International Business Machines Corporation, Armonk, NY, USA) after validation of normality with Shapiro-Wilk testing. A Greenhouse-Geisser correction for sphericity was used where appropriate. The full results of the statistical testing in SPSS are included in the GraphPad prism file available on the figshare repository. Post-hoc statistical testing versus the baseline state was performed with the Holm-Bonferroni correction for multiple comparisons.

Additionally, total vasomotion profile difference, a statistical group difference measure, was calculated using non-parametric cumulative mass-based statistical testing. In brief, the sum of the absolute value of the t-statistics thresholded at $|t| > 2.571$ computed from the occurrence histogram data for admittance gain, phase shift, and coherence for the INV and DNx kidneys were compared to those computed from the occurrence histogram data for shuffled null data, that is, occurrence histogram t-statistic maps derived from groups agnostic to denervation status. The cumulative thresholded t-statistic masses from each of the possible shuffled groups were calculated to create a non-parametric difference distribution. If a relationship between innervation status did not exist (null hypothesis), then the masses of these shuffled groups would be similar in magnitude to the actual INV vs. DNx data. On the other hand, if the mass of the actual data was greater than 95% of the cumulative differences for the shuffled groups, we rejected the null hypothesis at $\alpha = 0.05$ and concluded that there was a statistical relationship between innervation status and vasomotion.

**Power Analysis**

Prior preliminary data from three rabbits that underwent unilateral renal denervation indicated that the effect of denervation on measures of time-varying transfer function variability could be resolved with 80% power for an alpha = 0.05 for a sample size of 6 in the baseline state. While we hypothesized that hemorrhage would increase the magnitude of the observed effect, we did not know by how much the effect size would increase, and thus we decided to collect data from 6 rabbits.

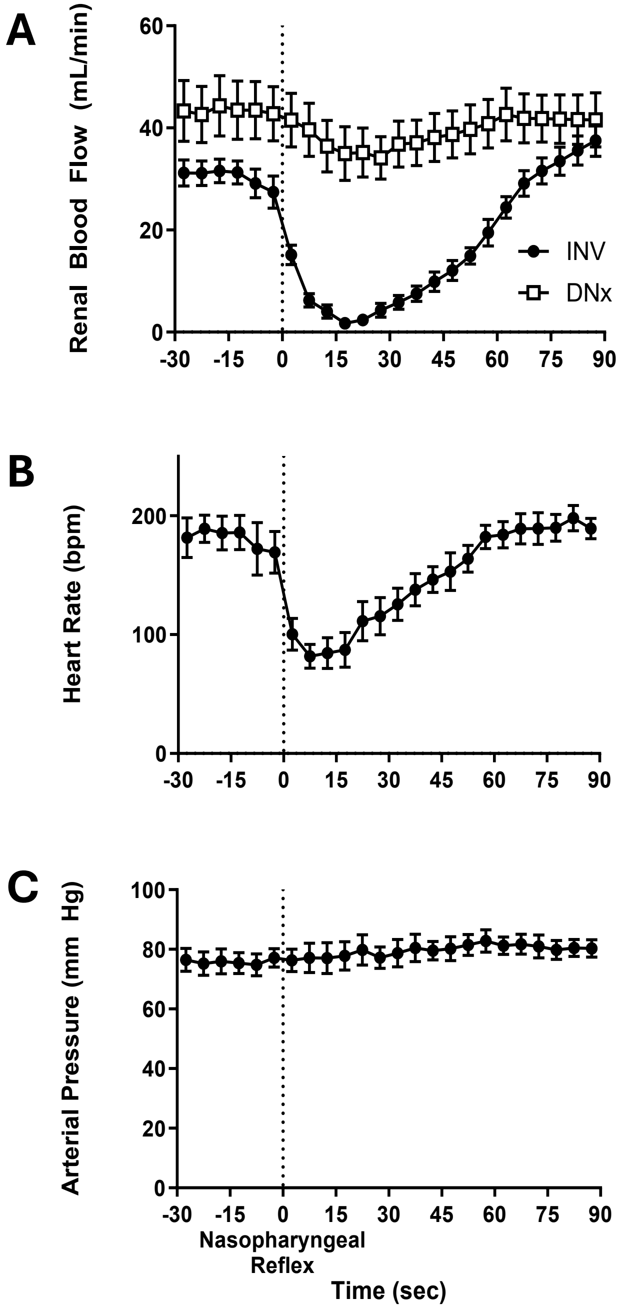


**Figure S1.** Nasopharyngeal reflex. Administration of a noxious respiratory stimulus to the rabbit results in a profound increase in sympathetic and parasympathetic outflow. This manifests as marked reductions in renal blood flow (A) in innervated kidneys that are abolished by renal denervation, providing a functional validation of the surgical denervation. This also produces profound decreases in heart rate (B). Despite profound opposing changes in autonomic outflow, arterial pressure (C) remains stable.

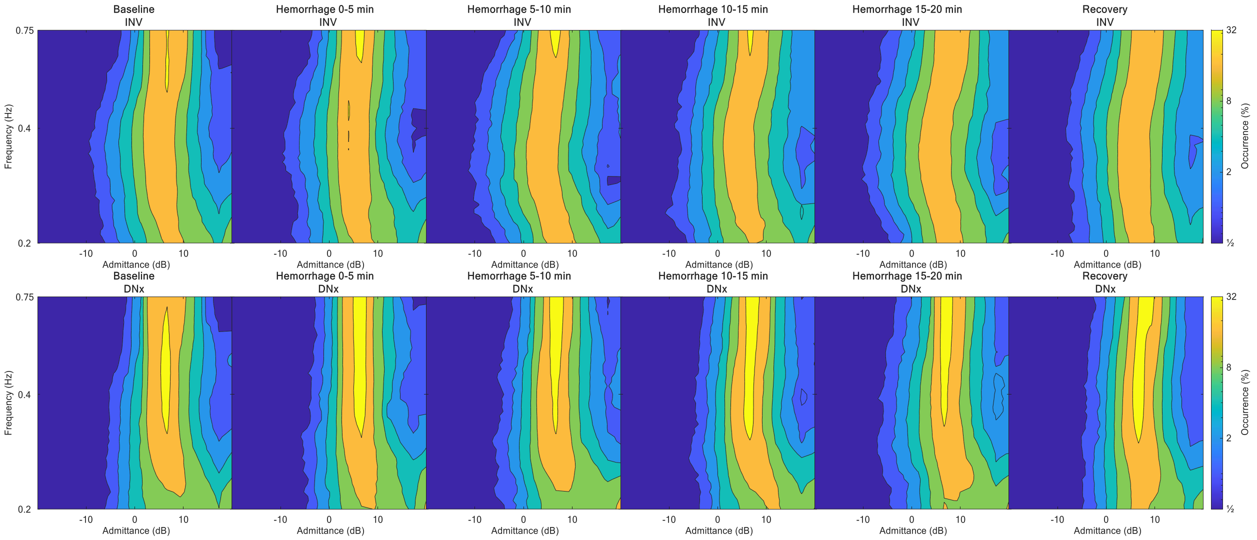


**Figure S2.** Admittance gain occurrence histograms. Mean admittance gain occurrence for INV (top) and DNx (bottom) kidneys shown as contour plots. Admittance gain is more consistent with more high admittance gain behavior in DNx kidneys while INV kidneys have more diverse admittance gain behavior with more frequent low admittance gain behavior. These differences are increased by hemorrhage.

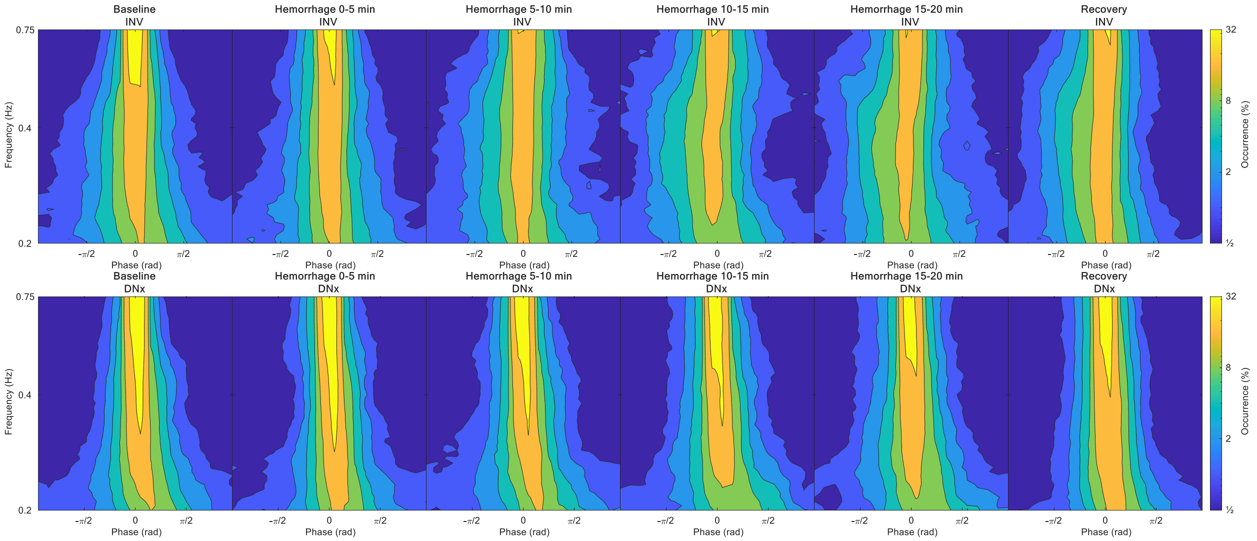


**Figure S3.** Phase shift occurrence histograms. Mean phase shift occurrence for INV (top) and DNx (bottom) kidneys shown as contour plots. Phase shift is more consistent with more zero phase shift (passive) behavior in DNx kidneys while INV kidneys exhibit more diverse and notably more frequent negative phase shift behavior, consistent with baroreflex-driven vascular control. These differences are increased by hemorrhage.

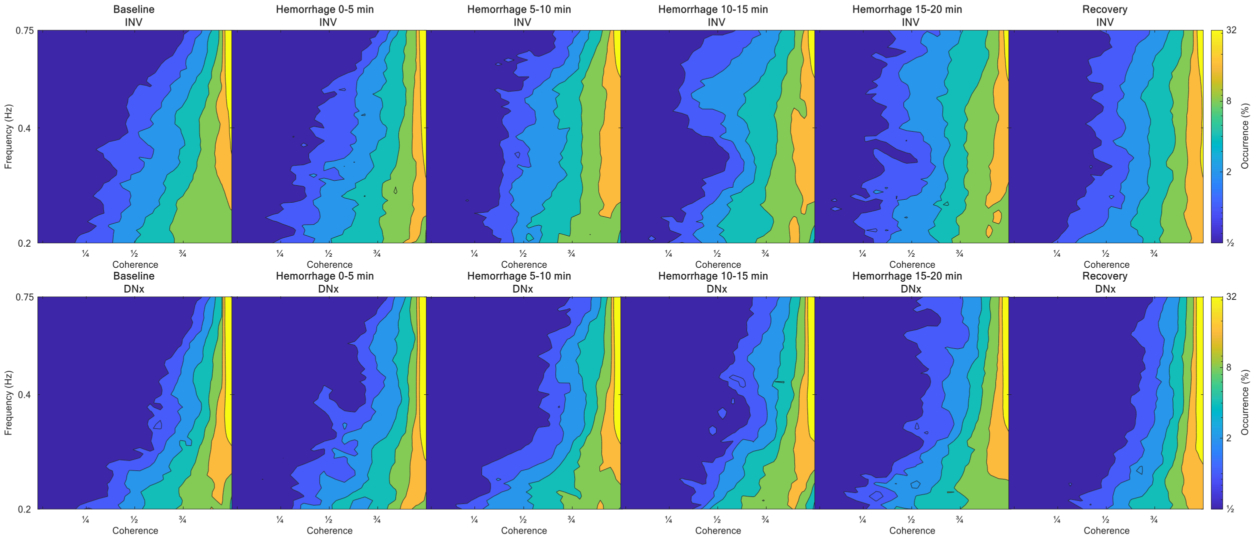


**Figure S4.** Coherence occurrence histograms. Mean coherence occurrence for INV (top) and DNx (bottom) kidneys shown as contour plots. Coherence is more consistent with more high coherence (passive) behavior in DNx kidneys while INV kidneys exhibit more diverse, low coherence behavior, consistent with active vascular control. These differences are increased by hemorrhage.
